# Identification of Barley Enhancers across Genome via STARR-seq

**DOI:** 10.1101/2022.12.10.519735

**Authors:** Wanlin Zhou, Haoran Shi, Zhiqiang Wang, Yuxin Huang, Lin Ni, Xudong Chen, Yan Liu, Haojie Li, Caixia Li, Yaxi Liu

**Affiliations:** State Key Laboratory of Crop Gene Exploration and Utilization in Southwest China, Chengdu 611130, China; Triticeae Research Institute, Sichuan Agricultural University, Chengdu 611130, China; Chengdu Academy of Agricultural and Forestry Sciences, Chengdu 611130, China

**Keywords:** Barley, Enhancer, Transposable elements, Gene expression, Functional validation

## Abstract

Enhancers are DNA sequences that can strengthen transcription initiation. However, the global identification of plant enhancers is complicated due to uncertainty in the distance and orientation of enhancers, especially in species with large genomes. In this study, we performed self-transcribing active regulatory region sequencing (STARR-seq) for the first time to identify enhancers across the barley genome. A total of 7323 enhancers were successfully identified, and among 45 randomly selected enhancers, over 75% were effective as validated by a dual-luciferase reporter assay system in the lower epidermis of tobacco leaves. Interestingly, up to 53.5% of the barley enhancers were repetitive sequences, especially transposable elements (TEs), thus reinforcing the vital role of repetitive enhancers in gene expression. Both the common active transcription marker H3K4me3 and repressive histone marker H3K27me3 were abundant among the barley STARR-seq enhancers. In addition, the functional range of barley STARR-seq enhancers seemed much broader than that of rice or maize and extended to ± 100 KB of the gene body, and this finding was consistent with the high expression levels of genes in the genome. This work specifically depicts the unique features of barley enhancers and provides available barley enhancers for further utilization.

## Introduction

Enhancers are among the most important *cis*-regulatory modules and help plants coordinate both developmental and environmental responses [1]. These common elements act in upstream or downstream of genes, and they are always located in distal promoter regions [2]. In recent years, a number of enhancers have been found to regulate developmental or tolerance processes, such as the inflorescence architecture, phosphate homeostasis, shoot regeneration, floral transition, and salinity stress tolerance [3–7]; thus, these elements show good potential in regulating various functional genes. However, enhancers are difficult to identify because their position and orientation are uncertain.

Multiple sequencing methods have been used to identify enhancers in plants. DNA and chromatin features have been used to predict enhancer candidates in both animal and plant genomes. DNase I hypersensitive sites (DHSs) are commonly associated with enhancers, silencers, promoters, insulators, and locus control regions [8]. However, the follow-up standard Southern blot approach is a complicated, time-consuming, and inaccurate process and thus represents a huge constraint in enhancer exploration [9]. To address this issue, massively parallel signature sequencing (MPSS) was combined with DHS mapping to promote genome-wide enhancer identification [9]. DHS data were recently obtained for species with relative small genomes, such as *Arabidopsis*, maize, tomato, sorghum, foxtail millet, and rice [10–12]. The assay for transposase-accessible chromatin with sequencing (ATAC-seq) method was recently introduced for the discovery of *cis*-regulatory region in plants [13]. However, the regulatory units identified by ATAC-seq are tissue-specific, thereby masking partial enhancers. DNA methylation and histone modification are considered important markers for identifying *cis*-regulatory units of grass genomes n[14–15]. In maize, DNA methylation has been successfully used to explore unmethylated regions (UMRs) that carry potentially active promoters and *cis*-regulatory units [15]. Thus, chromatin immunoprecipitation sequencing (ChIP-seq) technology has been utilized to discover enhancer sites with symbolic H3K4me1 and H3K27ac modifications in large quantities, thus offering the possibility of identifying super enhancers (SEs) [16–18]. Although the above methods can identify valid enhancers, only a limited number of enhancers have been found by DNA and RNA labels, and the current protocols are too expensive and labor intensive to apply to plant genomes.

Self-transcribing active regulatory region sequencing (STARR-seq), clustered regularly interspaced short palindromic repeats (CRISPR)-based approaches and massively parallel reporter assay (MPRA) are the three main methods of combining genomic identification with the functional utilization of enhancers, and STARR-seq is widely used to directly and quantitatively assess enhancer activity for millions of candidates from arbitrary sources of DNA in both mammalian and non-mammalian genomes [19–20]. Compared to other sequencing methods, STARR-seq can quantify enhancer strength in complex candidate libraries [21]. All enhancers, whether active, chromatin-masked, or dormant, can be identified through STARR-seq [22]. Enhancers in the genomes of *Drosophila*, humans, and mice have been successfully obtained based on the library construction protocol [19,22–24], and this method can provide a considerable number of candidates for further functional studies in animals. With the advancement of library construction by protoplast transfection, quantitative enhancers in *rice* and maize have also been discovered [25–26], and modifications of this method allow enhancers to be identified by STARR-seq and validated in transient tobacco systems [27]. Moreover, UMI-STARR-seq, a novel STARR-seq variant that includes unique molecular identifiers (UMIs), was developed to accurately count reporter mRNAs in low-complexity STARR-seq libraries, thus indicating that the accuracy of global enhancer identification is increasing [28].

Although *cis*-regulation by enhancers is vital, few enhancers have been validated in Triticeae crops, such as wheat and barley. In 1987, a 268-bp enhancer-like sequence was discovered in the 5’-proximal region of the wheat *chlorophyll a/b-binding-1* (*cab-1*) gene [29]; and in 1994, an enhancer/silencer sequence that directs aleurone-specific expression of a barley chitinase gene was identified [30]. Approximately three decades passed, only a few enhancers have been identified in barley and wheat due to the genome complexity and bulkiness of these plants. In this study, global enhancers in barley were identified for the first time using STARR-seq, and their strengths were then predicted and quantified. The specific sequence signature and ChIP-seq histone modification markers of barley enhancers, which are different from those of rice or maize, were also identified in this study.

## Results

### Total of 7323 barley enhancers were identified using STARR-seq

After STARR-seq, two replicates were merged to identify possible enhancers. In total, 398,293,684 reads were acquired in the input plasmid libraries, and the fragments had a median length of 518 bp; and 512,678,889 reads were obtained in the cDNA libraries, and the fragments had a median length of 535 bp. After eliminating duplicates, 98,972,877 reads were reserved in the plasmid libraries and 111,427,056 reads were reserved in the cDNA libraries.

In total, 7323 enriched peaks were successfully identified and used for follow-up analysis (**Figure 1A**; Table S1). Analysis of chromosomal locations indicated that 7323 STARR-seq barley enhancers were randomly located on seven chromosomes without obvious location preferences, indicating that our identification was genome wide (Figure 1B).

**Figure 1.**
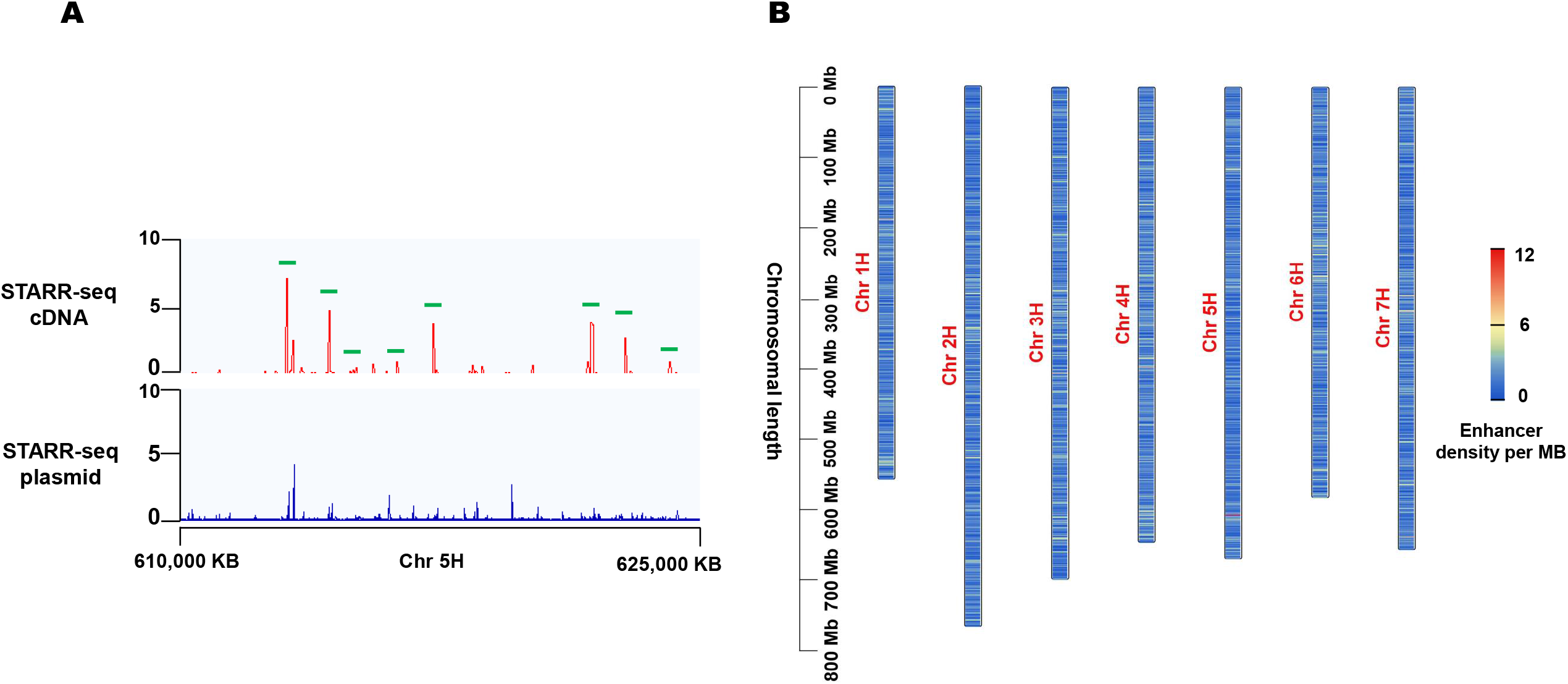
Genomic discovery of barley STARR-seq enhancers. **A**. Merged STARR-seq input plasmid (blue) and cDNA (red) reads at representative genomic region on chr 5H. Green boxes represented potential enhancer peaks. **B**. Distribution of STARR-seq barley enhancers across 7 chromosomes. Chromosomal length was labeled on the left and the scale bar was on the right. Scale values of the corresponding color represented the number of enhancers per MB (enhancer density). STARR-seq, self-transcribing active regulatory region sequencing.

### Few STARR-seq enhancers overlapped with ATAC-seq peaks in barley

Compared with previously published barley ATAC-seq peaks, our STARR-seq enhancers were more balanced in terms of distribution and density (Figure 1B and Figure S1). Interestingly, STARR-seq enhancers in rice were more balanced than ATAC-seq enhancers [25,31] (Figure S2). ATAC-seq peaks in rice, maize, and barley [31] were analyzed and found to be enriched at the ends of each chromosome, which is possibly because of bias in chromosome cleavage via transposase Tn5 (Figure S1, S2, and S3). The difference in distribution features further revealed that the STARR-seq method was less affected by chromosomal location or orientation. Meanwhile, barley ATAC-seq peaks were compared with our STARR-seq peaks for cross-validation. Surprisingly, few STARR-seq enhancers (∼1.3%) overlapped with ATAC-seq peaks in barley (Figure S1; Table S1). Similar to the poor intersection (∼8.7%) observed between ATAC-seq and DHS-seq enhancers in rice, the low proportion of overlapping enhancers among STARR-seq, ATAC-seq, and DHS-seq in plants was likely caused by different library construction principles [25].

### Low proportion of barley enhancers regulated adjacent target genes

Among the total barley STARR-seq enhancers, only 9%–11% in each chromosome directly overlapped with coding genes (**Figure 2A**). Furthermore, we searched for genes 10 KB upstream or downstream of the 7323 enhancers and found that a low proportion of enhancers (∼2735) had potential target genes at a distance of 10 KB (∼37%), which indicated that only a minority of genes in barley were regulated by immediately adjacent enhancers. However, in the rice genome, proximal regulation (± 5 KB) by enhancers may be more common because a proportion of over 70% was observed [25].

**Figure 2.**
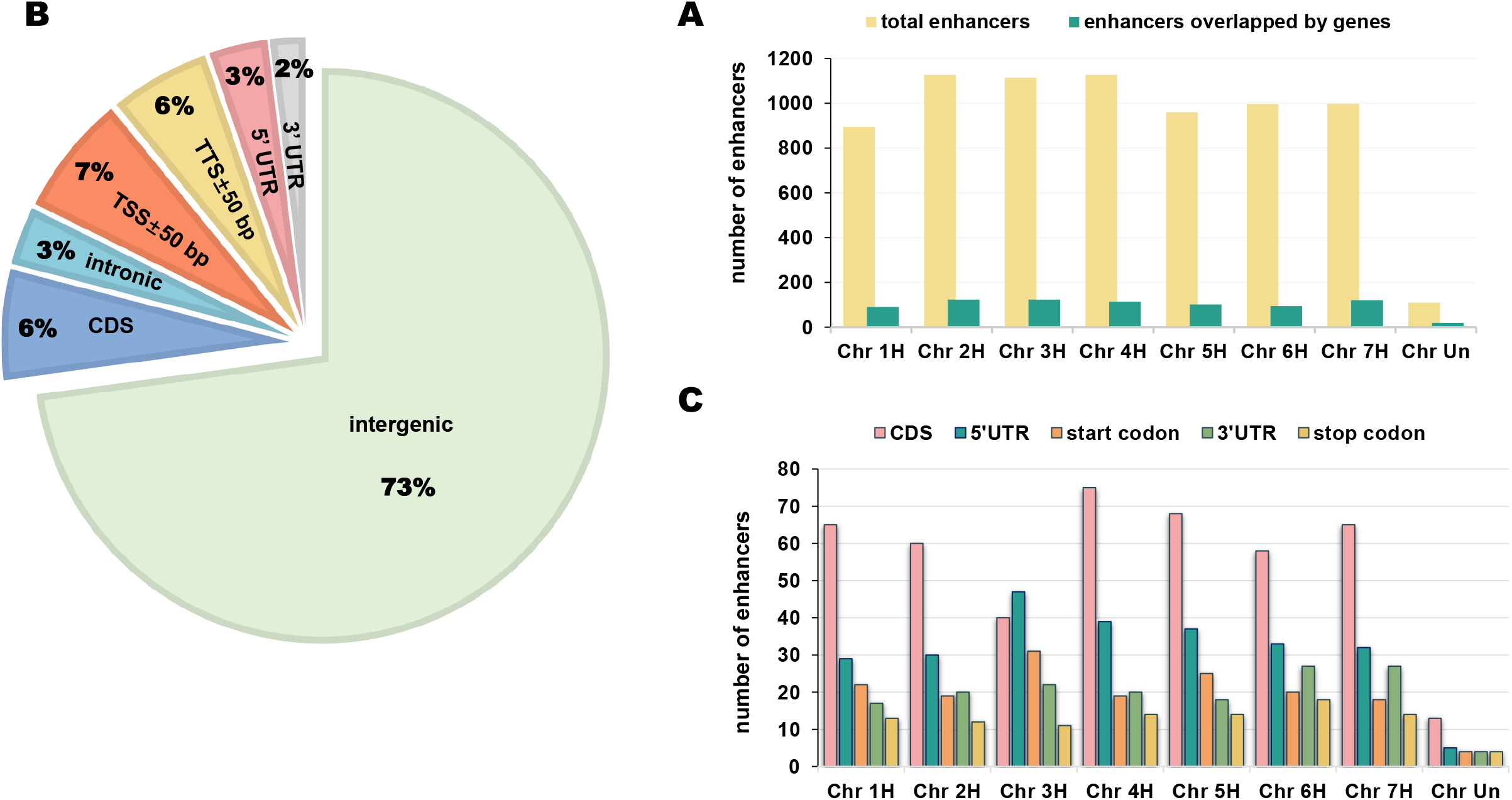
Position distribution characteristics of barley STARR-seq enhancers. **A**. Number of total enhancers and enhancers overlapped with genes in each chromosome. **B**. Proportions of enhancers overlapped with or without various gene elements. **C**. Number of enhancers overlapped with CDS, 5’UTR, start codon, 3’UTR, and stop codon regions. TSS, transcription start site; TTS, transcription termination site; CDS, coding sequence; 5’UTR, 5’ untranslated regions; 3’UTR, 3’ untranslated regions.

### Barley enhancers were enriched in intergenic regions

To determine the location preference of barley STARR-seq enhancers, all chromosomal locations of various gene motifs in barley, including intergenic regions, intronic regions, transcription start sites (TSSs), 5’ untranslated regions (5’UTRs), coding sequences (CDSs), 3’ untranslated regions (3’UTRs), and transcription termination sites (TTSs), were collected and blasted (Figure 2B).

Interestingly, barley enhancers covered intergenic regions (72.9 %) much more frequently than intronic (3.28%) and CDS regions (6.17%) (Figure 2B), and the distribution was quite different compared to that in the genomes of *Drosophila, Arabidopsis* and rice, which were smaller and less repetitive. In *Drosophila*, over 55% of the identified enhancers were located within introns, especially in the first intron (∼37.2%) [19]. In rice, less than 25% of enhancers mapped by STARR-seq were overrepresented in intergenic regions [25]. In *Arabidopsis*, 73.2% of STARR-seq enhancers were enriched in CDS regions, whereas only 0.2% of the enhancers were enriched in intergenic regions [32]. However, in highly repetitive maize genomes, approximately 70% of all mapped micrococcal nuclease hypersensitive site (MNase HS) sequences occurred within the intergenic space, despite different identification methods [33]. Thus, the distribution of STARR-seq enhancers between intergenic and non-intergenic regions might be influenced by the genomic size, transposable element (TE) composition, and gene density, regardless of the species.

Moreover, for genes overlapped by barley enhancers, the 5’UTR (∼3.28%) and TSS regions (∼6.79%) overlapped more than the 3’UTR (∼1.98%) and TTS regions (∼5.63%) (Figure 2C). Although the genomic size, TE composition, and intergenic spans of barley were quite different from those of rice and *Arabidopsis*, the enhancer distribution trend of 5’UTR > 3’UTR was consistent, thus revealing a common *cis*-regulation pattern in plants that enhancers prefer to locate at the 5’ end of genes [25,32]. This result also fits well with our universal understanding that enhancers always act in conjunction with promoter regions.

### Barley enhancers were highly repetitive in sequence

The enhancers identified in this study were aligned to the MorexV2.44_pseudomolecules_assembly genome, which revealed highly repetitive sequences. Repeatmasker identified 3917 enhancers (∼53.5%) with typical repetitive sequences of barley. These enhancers were further divided into three categories: class I retroposons, class II DNA transposons, and simple/tandem repetitive sequences, with approximately 2% of the enhancers involved in more than one repetitive category (**Figure 3A**). Class I retroposons accounted for the highest percentage (79.6%), and class II DNA transposons accounted for the lowest percentage (∼5.1%) (Figure 3B). The results revealed that a surprisingly high proportion of 84.7% STARR-seq enhancers in barley partially overlapped with TEs. In the rice genome, which presented a composition of approximately 35% TEs, 52.1% of the identified STARR-seq enhancers were located in the TE regions [25,34]. Obviously, STARR-seq enhancers within the TE regions of both rice and barley accounted for the majority, although the proportion seemed positively correlated with the genomic TE composition. Interestingly, in the maize genome, which is composed of 85% TEs, 4590 out of 32,421 (∼14.2%) ATAC-seq-identified accessible chromatin regions (ACRs) presented at least a partial overlap with annotated TEs [35]. Clearly, TEs generate enhancers to increase gene expression in cereal crops, although the proportion of TEs and enhancers is largely influenced by the sequencing method. Moreover, the results in cereal crops were quite different from that of *Drosophila* enhancers, which were significantly underrepresented in repetitive sequences [19,25].

**Figure 3.**
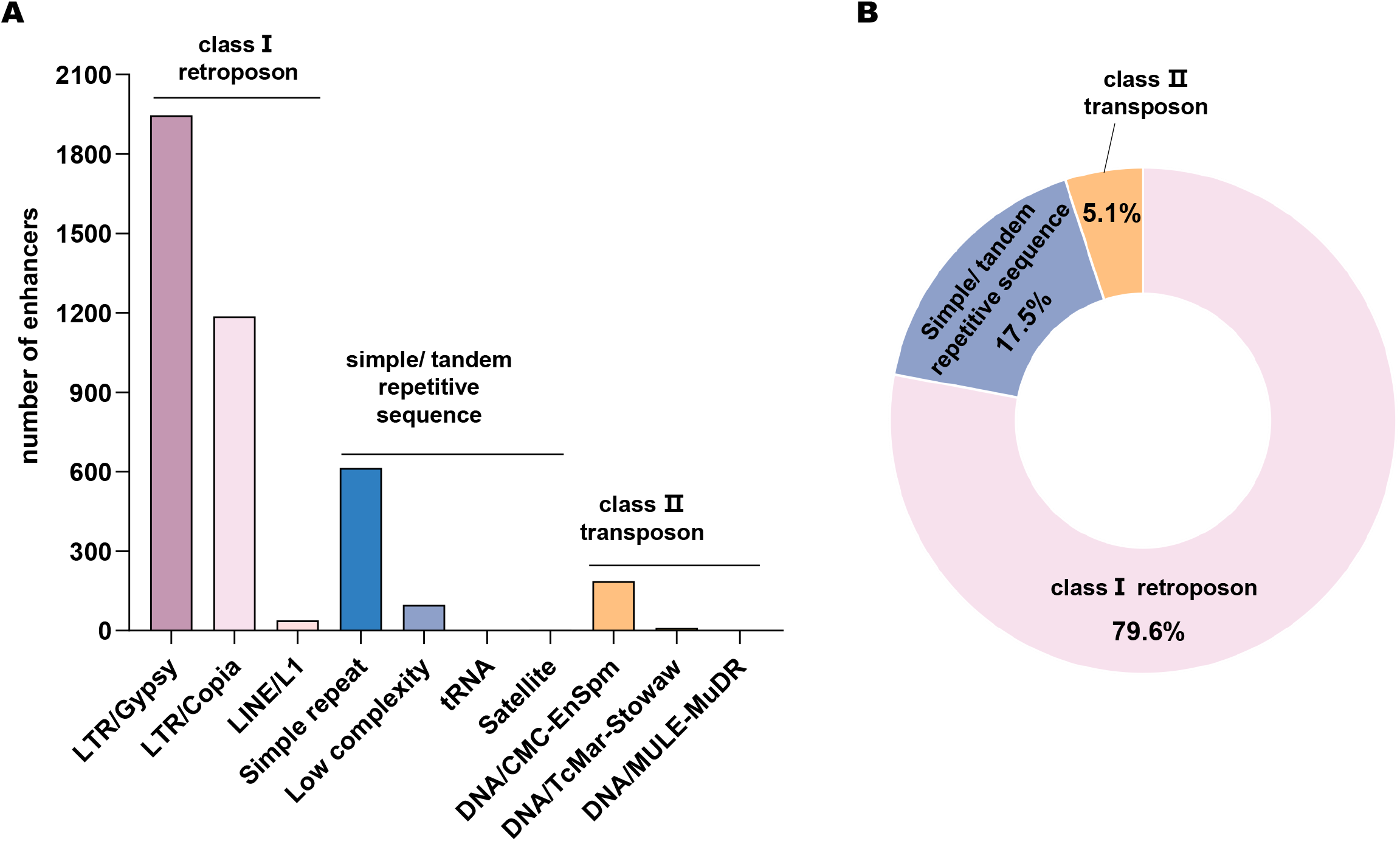
Sequence characteristics of barley STARR-seq enhancers. **A**. Number of enhancers in categories of various repetitive sequence. **B**. Proportions of enhancers in category of class [retroposon, class [DNA transposon and simple/tandem repetitive sequence, respectively. LTR, long terminal repeated; LINE, long interspersed nuclear elements; MULEs, Mu-like elements; Stowaw, a variant of miniature inverted repeat transposable elements.

More specifically, among the highly repetitive enhancers, 1946, 1186, and 39 enhancers were identified as long terminal repeat (LTR)/Gypsy, LTR/Copia, and long interspersed nuclear element (LINE)/L1 enhancers in the class □ retroposon category, respectively; and 614, 97, 3, and 1 enhancers were recognized as simple, low-complexity, satellite, and tRNA enhancers in the simple/tandem repetitive enhancer category, respectively; while 187, 10, and 2 enhancers were regarded as DNA/CMC-EnSpm, DNA/TcMar-Stowaw, and DNA/Mu-like element (MULEs)-MuDR enhancers among the class II DNA transposons, respectively (Figure 3A; Table S2). Analysis of the sequence signatures of highly repetitive enhancers may further reveal that LTRs, especially LTR/Gypsy, are significantly related to gene expression activities in barley, which is consistent with the fact that nearly 50% of barley TEs are LTR/Gypsy enhancers [36]. In maize, the LTR/Gypsy families were the most significantly enriched enhancer candidates [37]. In rice, LTR-retrotransposons could harbor regions with strong enhancer activities, and some STARR-seq rice enhancers indeed overlapped with LTRs, although the proportion was not clear [25,38]. TEs and LTRs are important components of plant enhancers, although in *Drosophila*, a similar sequence pattern was not observed.

### Barley enhancers may affect genes within 100 KB

In the rice genome, 28.7% of genes have at least one proximal enhancer (gene body ± 5 KB), with the majority of STARR-seq enhancers mapped within or close to genes [25]. In the larger maize genome, only 32.5% of ACRs occurred at >2 KB from their nearest genes and only 12.7% of enhancers occurred at >20 KB [26]. However, in barley genomes with significantly lower gene densities, the conclusions are different. Few genes had enhancers within the traditional ± 10 KB distance (∼5.8%). To roughly assess the functional range of barley enhancers, abundant RNA-seq data of barley genes were collected from a published online database, and then Fragments Per Kilobase Million (FPKM) values were calculated (Table S3). The overall expression levels of genes within 0–10, 10–50, 50–100, and ≥100 KB of STARR-seq enhancers indicated that genes located within 100 KB of barley enhancers presented higher expression relative to those located greater than 100 KB away, while the overall expression levels of genes within 0−10, 10−50, and 50−100 KB were similar (**Figure 4A**). This result suggests the possibility that barley enhancers influenced gene expression within a broader range of 100 KB. A large proportion of repetitive sequences and greater distances between genes in barley possibly blocked the enhancers from regulating nearby genes. Compared to rice (approximately 125 genes/MB) [34] and maize (approximately 20 genes/MB) [39], the lower gene density in barley (approximately 0.5 gene/MB) [40] might have driven enhancers to adapt to a much broader functional range.

**Figure 4.**
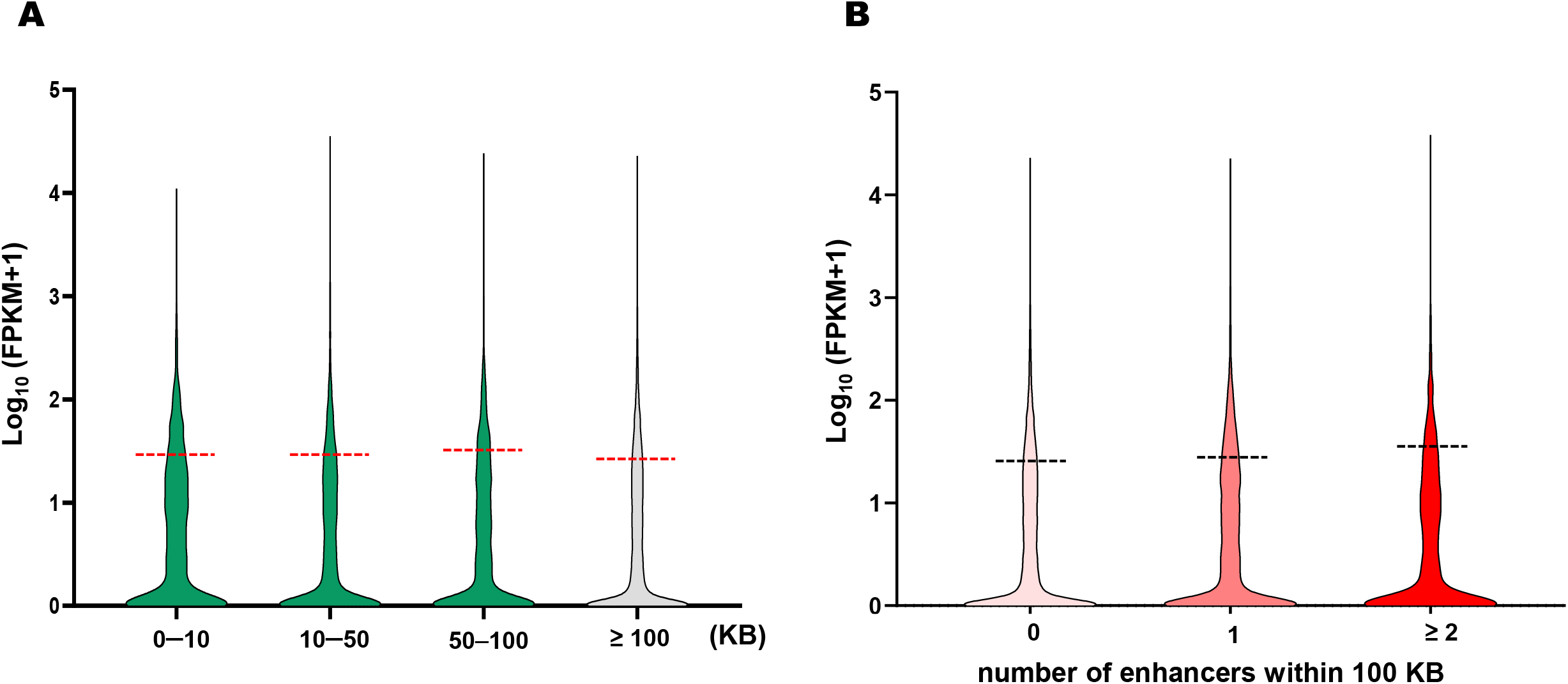
Functional range and enhancing effects of barley STARR-seq enhancers. **A**. Overall expression levels of genes within 0−10, 10−50, 50−100, and ≥ 100 KB of barley STARR-seq enhancers. **B**. Overall expression levels of genes having 0, 1, and ≥ 2 enhancers within 100 KB. FPKM, Fragments Per Kilobase Million.

To further validate this hypothesis, quantitative analyses of barley genes with or without enhancers in a 100 KB range were performed (Table S4). As expected, despite not reaching a significant level, the overall expression level of the 12,129 barley genes with enhancers within 100 KB was higher than that of the 30,702 barley genes without enhancers within 100 KB (average FPKM value: 28.95>26.5). More interestingly, the overall expression levels of genes with ≥2 enhancers within 100 KB were improved compared to those with only one enhancer (average FPKM value: 32.37>28.95), thus revealing potential accumulative effects of barley enhancers (Figure 4B). In general, the enhancers we identified worked as up-regulatory elements in barley at the omics level.

### Enhancers of different repeat categories played different roles in gene expression

Although class I retroposon TEs, namely, LTR/Copia and LTR/Gypsy enhancers, were identified as the main types of repetitive barley enhancers, their enhancing effects might not be the best (**Figure 5A**; Table S5). After separately analyzing the average expression levels of genes within 100 KB of enhancers in the no-repetitive-sequence category and 10 repetitive sequence categories (satellite, tRNA, LINE/L1, LTR/Copia, LTR/Gypsy, DNA/CMC-EnSpm, DNA/MULE-MuDR, DNA/TcMar-Stowaw, simple repeat, and low-complexity enhancers), enhancing effects were preliminarily assessed. Surprisingly, satellite, simple repeat, and low-complexity enhancers with the simplest tandem repeat sequences exhibited the best enhancing effects (Figure 5A; Table S5). Retroposon-related enhancers (LINE/L1, LTR/Copia, and LTR/Gypsy) could be regarded as tracing markers in the largest proportion, although their enhancing effects were not very strong (Figure 5A). Enhancers without repetitive sequences tended to play the weakest regulatory roles, thus revealing the importance of redundant repeats in barley *cis*-regulation (Figure 5A).

**Figure 5.**
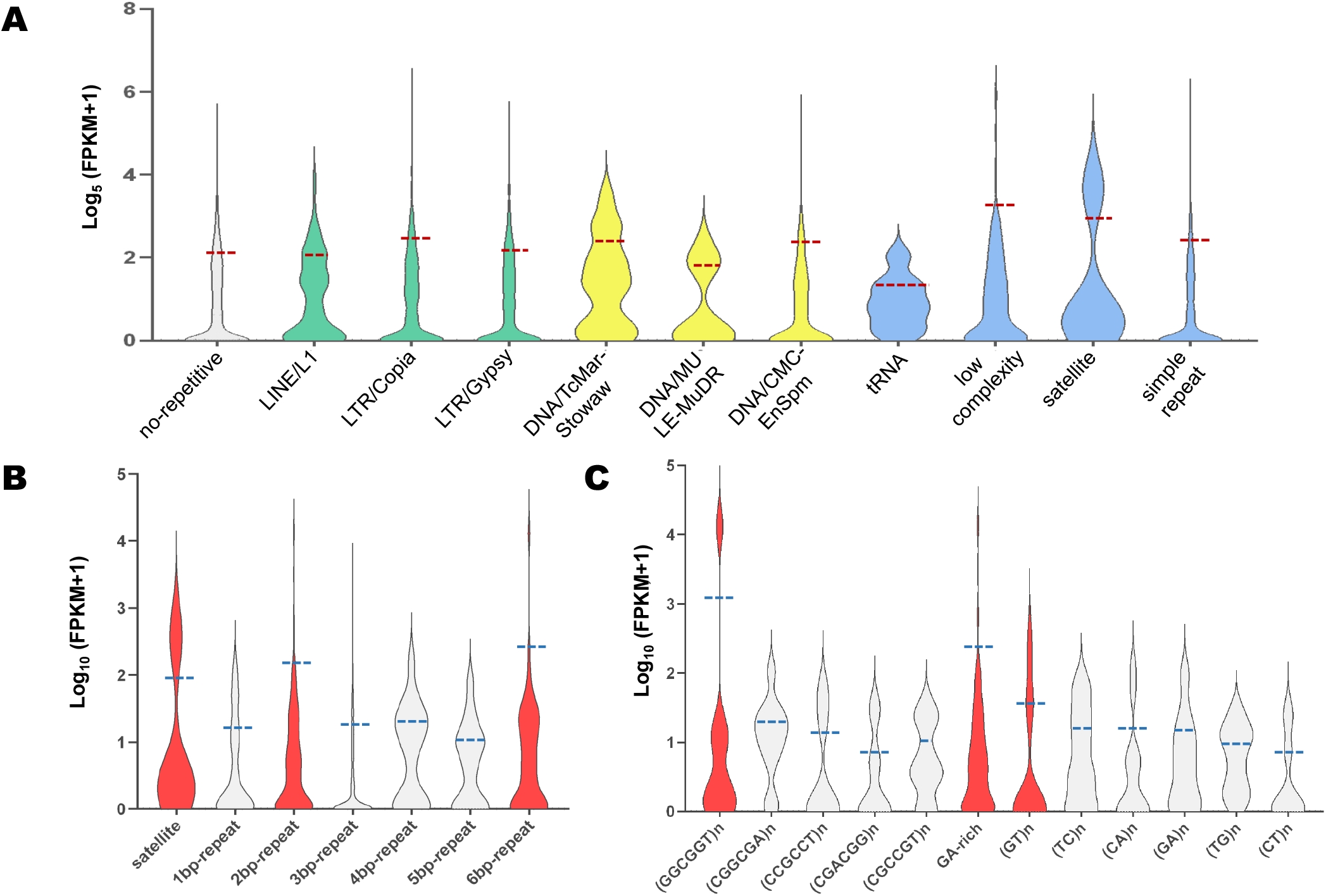
Enhancing effects analysis of barley STARR-seq enhancers in different categories of repetitive sequence. **A**. Overall expression levels of genes having no-repetitive sequence enhancers or having 10 categories of repetitive sequence enhancers. **B**. Overall expression levels of genes within 100 KB of enhancers in simple/tandem repetitive sequence category; the groups were divided by base pair number. **C**. Overall expression levels of genes within 100 KB of enhancers in 2bp-repeat and 6-bp repeat groups of (**B**); the groups were divided by base pair composition.

Moreover, more specific repetitive types of the best-performing satellite, simple repeat, and low-complexity enhancers were analyzed in detail. According to the number of base pairs, satellite, 2-base-pair-repeat, and 6-base-pair-repeat enhancers were the strongest types (Figure 5B; Table S6). Among all base compositions, GA-rich, (GT)n, (GGCGGT)n, or satellite enhancers had the most positive effect on gene expression levels within 100 KB at the genomic level (Figure 5C; Table S6).

### STARR-seq enhancers were successfully validated by a tobacco system

A total of 45 peaks were successfully cloned into the modified pGreen II-0800-*Luc* vector. Independent of the *cab-1* enhancer [29] and the empty vector as positive and negative controls, respectively, the dual-luciferase reporter assay system revealed that the majority of our enhancers were effective in promoting *luciferase* (reporter gene) expression via preliminary imaging. Randomly chosen enhancers with strong, medium, and weak enrichment values showed an obvious enhancing effect compared to the minimal 35S promoter only; however, the degrees were quite different. The partial chemi-imaging results are shown in **Figure 6**.

**Figure 6.**
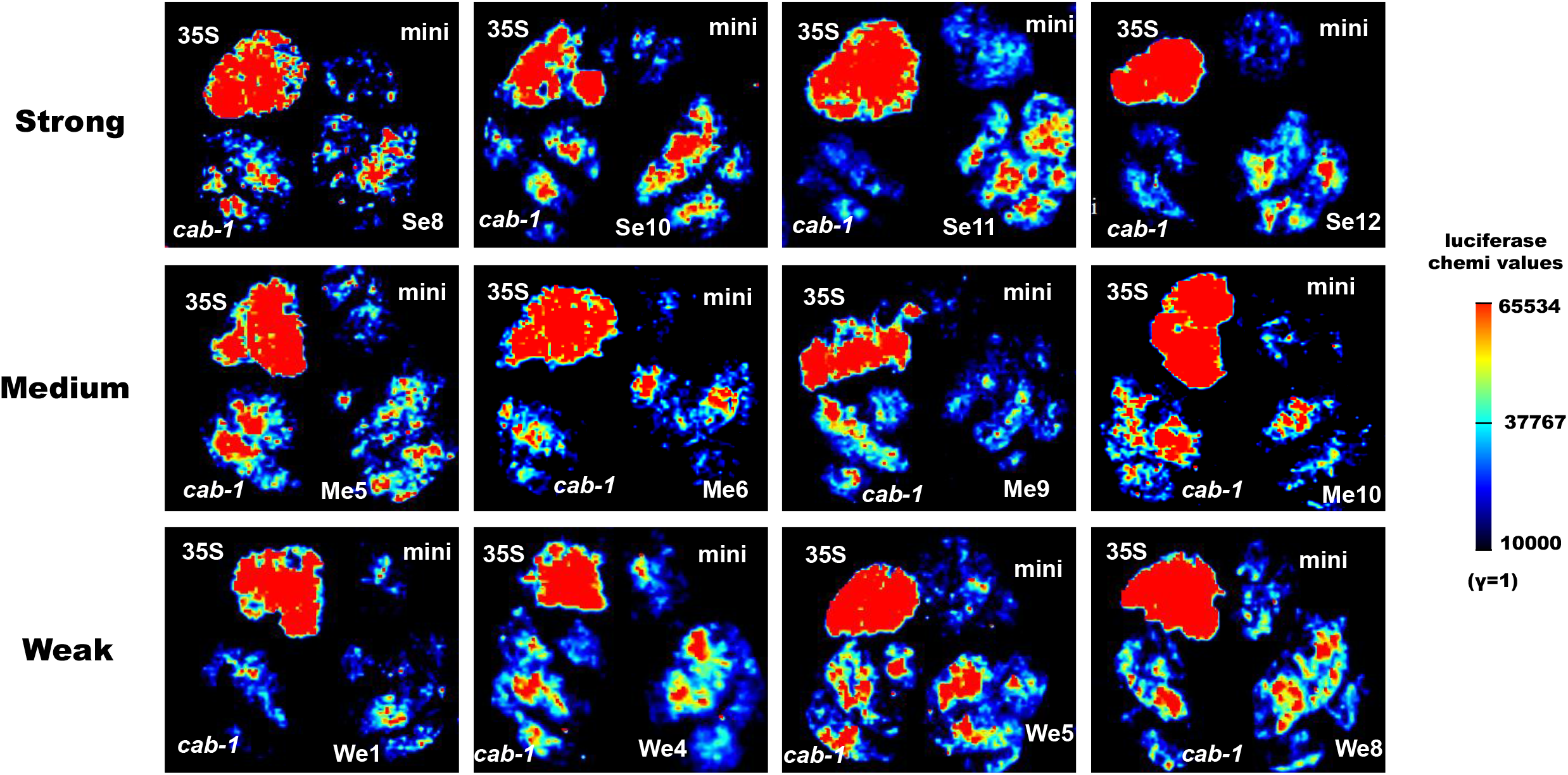
Chemical luciferase imaging of representative strong, medium, and weak enhancers. Enhancer *cab-1* was used as a positive control; 35S represented empty carrier with CaMV 35S promoter and was another positive control; mini represented empty carrier with minimal CaMV 35S promoter and was a negative control. Scale bar of luciferase chemical values referred to the “Image Lab” instrument parameter (γ=1). Se, Strong enhancer; Me, Medium enhancer; We, Weak enhancer.

To further quantify the effects of enhancers predicted by STARR-seq, quantitative reverse transcription PCR (RT-qPCR) was performed on the 45 enhancer treatments. As expected, 35 of 45 (∼77.8%) enhancer sequences were successfully validated, with 11, 15, and 9 sites having strong, medium, and weak strengths, respectively. The average enhanced levels were quantified by the enhancer/mini values (*luciferase* expression level of the minimal 35S promoter only was normalized to 1). The predicted strong enhancer sites increased the *luciferase* expression levels by an average of 1.885 times, which was higher than that of the medium sites (∼1.534 times) and weak sites (∼1.154 times) (**Figure 7A**). Notably, enhancer Se4 in the strong catalog even exceeded the effect of the reported positive control *cab-1*. Our results showed that the effects of enhancers in plants were highly correlated with our STARR-seq enrichment values (Figure 7B).

**Figure 7.**
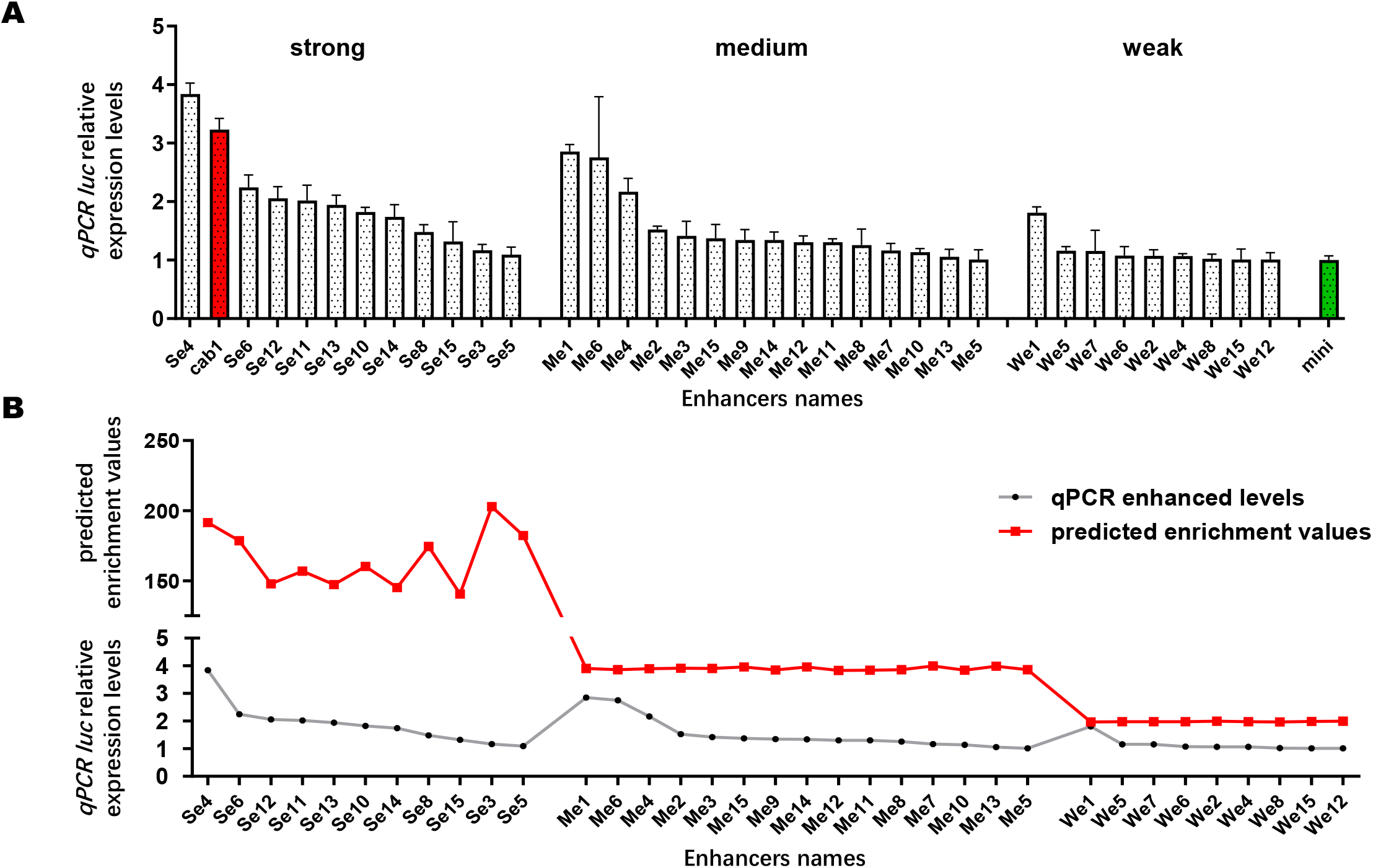
RT-qPCR validation of 35 effective STARR-seq enhancers. **A**. Relative expression levels of *luc* enhanced by 35 effective STARR-seq enhancers. 35 enhancers were included in the 45 randomly chosen enhancers from strong, medium, and weak categories. Expression level of *Renilla luciferase* was used as an internal control. Column in red represented the relative expression level of *luc* up-regulated by *cab-1* enhancer (positive control); column in green represented the relative expression level of *luc* promoted by minimal CaMV 35S only and was normalized to 1 (negative control). **B**. Fitted curves between predicted enrichment values and qPCR relative expression levels of *luc* enhanced by 35 effective enhancers. *Luc, Firefly luciferase*.

In conclusion, the enrichment levels of candidate enhancers were significantly associated with the upregulated levels of *luciferase*. Both the enhancer sequences and enhancing strength predicted from our STARR-seq data were reliable for further functional analysis and utilization.

### H3K4me3 and H3K27me3 were enriched at STARR-seq enhancers

Histone modification markers of barley STARR-seq enhancers were analyzed by collecting published ChIP-seq data. The active transcription marker H3K4me3 and repressive histone marker H3K27me3 were included in the analysis [41,42]. In a previous study, H3K4me3 and H3K27me3 were both enriched in rice STARR-seq enhancers [25]. A similar result was observed for histone modifications in barley.

Both the active transcription marker H3K4me3 and repressive histone marker H3K27me3 were enriched in our barley STARR-seq enhancers, with obvious peaks (**Figure 8**). Interestingly, H3K27me3 has been reported to restrict or inhibit enhancer activity [43–45]. Anomalous H3K27me3 enrichment, both in rice and barley, further confirmed that H3K27me3 is mostly associated with a repressed chromatin state in animals, although in plant genomes, it may represent a dependent enhancer marker regardless of whether it is latent or active.

**Figure 8.**
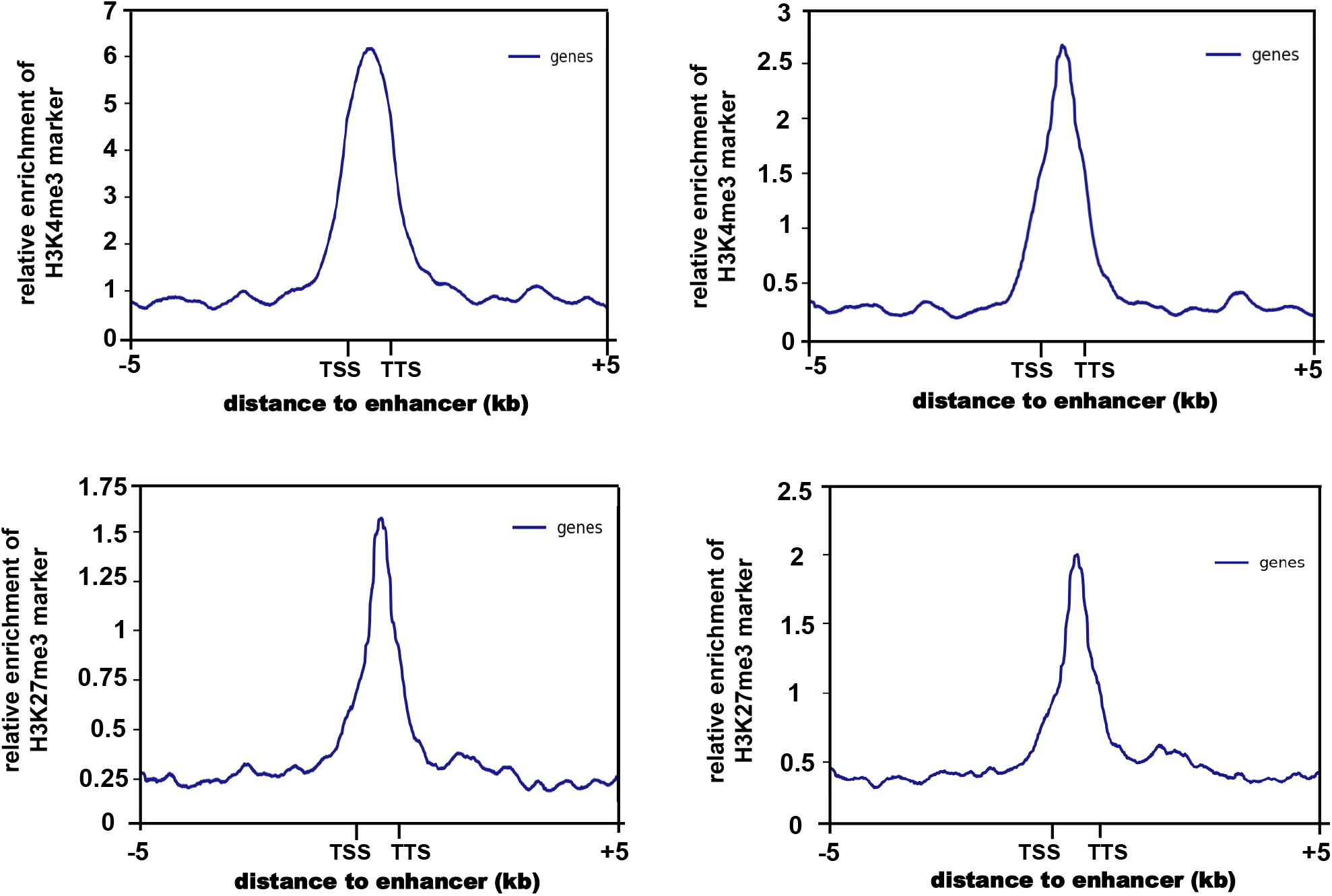
Histone modification markers enriched at barley STARR-seq enhancers. ChIP-seq data of H3K4me3 and H3K27me3 markers within 5 kb of STARR-seq enhancers were collected. The distance from TSS to TTS represented the enhancer body; the blue peaks represented the enrichment.

Throughout the genomes of mice, *Drosophila*, humans, rice, maize, and wheat, H3K27ac, H3K9ac, and H3K36me3 are conserved enhancer markers while H3K4me1 is underrepresented in rice, maize, and wheat but actively enriched in *Drosophila* and humans [15,19,25,37,46,47]. Unfortunately, ChIP-seq data have not been reported on H3K27ac, H3K9ac, H3K36me3, or H3K4me1 in barley.

## Discussion

### Barley enhancers identified by STARR-seq were highly reliable

The expression levels of barley mRNAs with or without STARR-seq enhancers within 100 KB revealed the authenticity and reliability of the 7323 predicted enhancers at the omics level, and mRNAs with STARR-seq enhancers within 100 KB tended to exhibit significantly higher expression. Moreover, the dual-luciferase reporter assay system experimentally illustrated the reliability of our predicted enhancers, with over 70% of the candidates validated in heterologous systems. However, because certain tissue-specific enhancers are stimulated by light or time, the system itself cannot fully represent the potential of our enhancers with hidden functions [27,48]. The remaining enhancers that do not work in tobacco may also work well in barley. Imaging and qPCR results in tobacco revealed the authenticity and reliability of 7323 predicted enhancers at the experimental level.

### Commonalities and characteristics of barley enhancers

The location signature of barley enhancers was further investigated in this study. STARR-seq enhancers identified in the *Drosophila, Arabidopsis* and rice genomes were mostly located close to or within genes [19,25,32]. Among the maize and barley enhancers, the majority were located in intergenic regions, which was partially because of the different genomic compositions [33]. Interestingly, compared to the 3’UTR and TTS ± 50 bp regions, the barley, rice, and *Arabidopsis* STARR-seq enhancers had a relatively higher proportion in the 5’UTR and TSS ± 50 bp regions [25,32]. The results mirrored the special location preference of plant enhancers. The activity of enhancers is always lower when located downstream of the gene than upstream of the minimal 35S promoter [27]. The location distribution feature may help maximally utilize plant enhancers.

Meanwhile, the sequence signatures of plant enhancers tend to be highly repetitive in the grass family, with the majority of enhancers existing as TEs [25,35,49–51]. Despite their commonality, barley STARR-seq enhancers also have unique sequence features. In barley, a significantly greater number of retroposon-related enhancers (∼79.6%), especially LTR/Gypsy enhancers, was observed relative to DNA transposon-related enhancers (5.1%). An evolutionary analysis among Triticeae crops indicated that the LTR/Gypsy element was one of the main reasons for genome expansion [36]. High-proportion LTR/Gypsy enhancers might play a vital role in both gene expression and evolution, and retroposons could represent an available label for tracing enhancers in both animals and plants [52,53].

A broader functional range of barley enhancers has also been proposed. Traditionally, 10 KB or less has been regarded as the strengthening range of enhancers in both mammals and plants with small genomes, such as rice and *Arabidopsis* [6,53–56]. However, barley enhancers have the potential to significantly broaden the range to 100 KB via gene expression analysis. As reported in maize, the long interaction enhancers *booster1* (*b1*), *teosinte branched1* (*tb1*), *vegetative to generative transition 1* (*Vgt1*), and *dIstal cis-element* (*DICE*) could *cis-*regulate target genes at distances of ∼100, 70, 60, or even 140 KB [1,3,57–59]. The high proportion of TEs and large intergenic spaces in barley not only generate enhancers but might also promote their regulation of additional genes.

### Repetitive enhancers have huge potential for gene regulation

The role of repetitive sequences is further emphasized in this study, and LTRs have long been reported to enhance human responses [60–62]. In plants, an 853 bp tandem repeat sequence located 100 KB upstream of *B1* regulates *B1* expression in maize [57,63]. A miniature inverted-repeat TE inserted 70 KB upstream of the maize *AP2/ERF* transcription factor (*ZmRap2*.*7*) could regulate *ZmRAP2*.*7* expression by affecting epigenetic modification [58]. In this study, the expression analysis indicated that enhancers of simple/tandem repetitive sequences had the best enhancing activity, thereby partially contradicting the previous hypothesis that tandem repeats acted as weak enhancer silencers to modulate the expression of proximal genes [64]. Our study further emphasizes the importance of enhancers in repetitive sequences and indicate that they might strongly enhance the expression levels of important functional genes to adapt to a transient environment.

In addition, three dominant repeat types of barley enhancers, GA-rich, (GT)n, and (GGCGGT)n, were identified. Tandemly repeated GA-rich sequences were frequently reported as motifs within highly active promoter regions and thus showed good potential as strong enhancers in animals and humans [65–67]. A (GT)n polymorphism in the promoter/enhancer region of *forkhead box protein P3* (*FOXP3*) is associated with the development of severe acute graft-versus-host disease (GVHD) in humans [68]. Although such effects have not been reported in plants, in our study, the RNA-seq data analysis partially supported the enhancing effects of “GA-rich” and “(GT)n” enhancers. These two enhancers may be common and conserved sequence signatures of the active enhancers. In addition, the newly discovered (GGCGGT)n enhancer may be another sequence signature for tracing highly active plant enhancers.

## Conclusion

In summary, we identified 7323 enhancers in barley and validated their authenticity via RNA sequencing data analysis and experimental RT-qPCR quantification.Intergenic and repetitive barley enhancers were found in high proportions, and the markers H3K4me3 and H3K27me3 were enriched. LTR enhancers were the most enriched, while simple tandem repetitive enhancers showed the greatest enhancement, thus demonstrating enhancer characteristics unique to barley.

## Materials and methods

### Reporter plasmid construction and library preparation

To comprehensively identify sequences with enhancer activity in the barley cultivar Morex, we constructed reporter and plasmid libraries from randomly ultrasonicated genomic DNA ranging from 500 to 800 bp. The input plasmid DNA library was separately transfected into protoplasts multiple times (4−6 times) to increase the abundance, and protoplasts were transfected and then randomly divided into two replicates for library construction.

The library preparation protocols mainly referred to a previous study on rice with the same modified plasmid pBI221 [25]. First, 1 ml genomic DNA (>100 ng/μL) was obtained from two-week-old barley seedlings. Genomic DNA was then fragmented by non-contact sonication (20% power at 5 s on and 10 s off, which was repeated 14 times in a liquid volume of 1 ml). DNA fragments ranging from 500 to 800 bp were separated by 1% agarose electrophoresis and maximally collected using a FastPure^®^ Gel DNA Extraction Mini Kit (Catalog No. DC301-01, Vazyme Biotech, China). The purified DNA products were repaired and ligated to VAHTS™ DNA Adapters Set 1/Set 2 for Illumina® (Catalog No. N801/802-01/02, Vazyme Biotech). The ClonExprressII One-Step Cloning Kit (Catalog No. C112, Vazyme Biotech) was used to successfully ligate the adaptor-ligated genomic DNA into the pBI221 vector, which was linearized by the incision enzymes BsrG□ and Mlu□. Different ratios (>10) of DNA fragments/linearized pBI221 vectors were generated to avoid connection preference of short fragments. Ligation products were transformed into DH5α strains (Catalog No. DL1001; Weidi Biotechnology, China) and cultured at 37□ for 12−16 h. The cultivation volumes were further increased to 5 to 6 L using Luria-Bertani medium, and the reporter plasmids were purified using the E.Z.N.A.^®^ Endo-Free Plasmid Maxi Kit (Catalog No. D6926, Omega Bio-tek, Norcross, GA). The bacterial endotoxin of the input plasmid was removed by adding 5 mol/L NaCl, a mixture solution of 30% polyethylene glycol 6000 (PEG 6000) + 1.5 mol/L NaCl, and 70% ethanol. The purified plasmid was collected and quantified using a NanoDrop ONE (Thermo Fisher Scientific, Waltham, MA).

### Protoplast transfection

Protoplasts were isolated from 20 to 30 etiolated leaves of barley seedling leaves after 2 weeks of cultivation. Etiolated seedlings were chopped into 0.5−1 mm strips and digested with cellulase R-10 (Catalog No. 9012-54-8, BEIJING LABLEAD BIOTECH, China) and Macerozyme® R-10 (Catalog No. 200115-03, Yakult Pharmaceutical Inc., Japan). The cell walls were digested for approximately 4 h by shaking at 45−50 rpm under dark conditions at 25 °C (ZWY-100H, China). A 200 nylon mesh filter was used to sieve the enzymatic hydrolysate, and W5 solution (154 mM NaCl, 125 mM CaCl_2_, 5 mM KCl, and 2 mM MES, pH 5.7) was used to repeatedly wash the protoplasts. Protoplasts were collected by centrifugation at 4 °C and 100 × g. For transfection, 40 μg of reporter plasmid DNA (≥ 2 μg/μL) was gently blended with 200 μl of protoplasts (approximately 1×10^6^ cells) in a non-stick centrifuge tube, and then 220 μl of freshly prepared PEG 4000 solution (40%, w/v) (Catalog No. 807490, Sigma-Aldrich Biotech, St. Louis, MO) was added to mediate transfection in the dark. The W5 solution was used again to wash off the 40% PEG and terminate the reaction after 15−20 min. Finally, the protoplasts were incubated in W5 solution at 25 □ for 16 h under dark conditions.

### Illumina sequencing library construction for reporter cDNAs and input plasmids

A VAHTS™ Universal DNA Library Prep Kit for Illumina^®^ V3 (catalog no. ND607, Vazyme Biotech) and VAHTS™ mRNA-seq V2 Library Prep Kit for Illumina^®^ (Catalog No. NR601, Vazyme Biotech) were used to construct input plasmid libraries and reporter cDNA libraries, respectively, using nearly the same flow path. Specifically, both mRNA and plasmid DNA from the transfected protoplasts were extracted using a Plant RNA Collection Kit v1.5 (Catalog No. RN33050, Biofit, China) without DNA elimination. VAHTS mRNA Capture Beads (Catalog No. N401, Vazyme Biotech) were used to capture RNA only, and VAHTS DNA Clean Beads (Catalog No. N411; Vazyme Biotech) were used to capture DNA from the remaining solution. cDNA was obtained using PrimeScript Reverse Transcriptase (Catalog No. 2690A, Takara, Japan) using a DNA elimination process.

Next, nested PCR was performed to amplify the cDNA and DNA using Phanta Max Super-Fidelity DNA Polymerase (Catalog No. P505; Vazyme Biotech) for less than 25 cycles in total. PCR products from the first round were purified using VAHTS DNA Clean Beads (0.8×) and then used as templates for the second round of PCR amplification with primers, including VAHTS™ DNA Adapters for Illumina^®^. VAHTS DNA Clean Beads (0.8×) were used again for purification. The total PCR cycles of the cDNA and DNA libraries should be the same and as few as possible. Both libraries were sequenced on a NovaSeq-150PE platform at Berry Genomics (Beijing, China).

### Identification of barley STARR-seq enhancers

The identification procedure for STARR-seq data was based on Sun et al. [25] with some adjustments. First, the raw reads of the two replicates were merged to maximize the STARR-seq enhancers. Bowtie2 was used to map the sequence data to the *Hordeum vulgare* genome (Morex V2.44 pseudomolecules; http://plants.ensembl.org/Hordeum_vulgare/Info/Index) [69,70]. SAMtools was used to filter the mapped reads and store read alignments against the reference sequences [71]. Redundant data caused by PCR amplification was eliminated using the “default” parameters [72]. The R package was used for STARR-seq enhancer identification [19]. BasicSTARRseq [73] and Bonferroni correction [74] were performed to adjust the *P* values. The genomic region was identified as an enhancer “peak” if the enrichment of cDNA over the input plasmid insert was >1.3 fold and the adjusted *P* value was <0.001.

### Location and sequence characteristic analysis of identified enhancers

Chromosomal information, location information, and gene annotation files were downloaded from the MorexV2.44_pseudomolecules_assembly reference genome of ***Ensembl Plants*** (http://plants.ensembl.org/index.html). Based on the above information, the locations of all identified STARR-seq enhancers were subjected to an orderly blast search with various gene elements to obtain position relationships and preferences. The distribution of enhancers across the barley genome was depicted by the Gene Location Visualize function in TBtools [75]. RepeatMasker was used to blast 7323 enhancers with known repetitive types in barley [76], and detailed repetitive sequence names and categories were output directly.

### Comparison between STARR-seq enhancers and ATAC-seq enhancers

ATAC-seq data for barley were collected from a previous study to validate the STARR-seq enhancers [13]. STARR-seq enhancers whose peak locations fell into the peak ranges of the ATAC-seq data were considered “overlapped”. With the help of the Gene Location Visualize function in TBtools, the chromosomal distribution and density characteristics of the STARR-seq enhancers in rice and barley and ATAC-seq peaks in rice, barley and maize were all determined [75]. Data on ATAC-seq peaks in barley, maize, and rice and data from STARR-seq enhancers in rice were collected from previously published supplementary materials [13,25].

### Expression analysis of enhancers and genes in barley

Approximately 42,831 genes were annotated in the whole barley genome of nearly 5,100,000,000 bp, indicating that the average distribution of barley genes was approximately 1 gene/100 KB. To assess whether the functional range of barley enhancers was within 100 KB, the overall expression levels of genes within 0–10, 10–50, 50–100, and ≥100 KB of STARR-seq enhancers were calculated. The RNA-seq data of 228 samples that were randomly related to abiotic resistance, disease resistance, and growth and development were collected from published online databases in the National Center for Biotechnology Information (NCBI) (accession numbers: PRJNA431836, PRJNA495764, PRJNA496380, PRJNA558196, PRJEB39864, PRJEB34186, GSE167271, PRJNA639036) [77–84]. RSEM software was used to quantify all genes, and the FPKM values were acquired [85]. To further check the 100 KB functional range, FPKM values of genes without enhancers within 100 KB, genes with one enhancer within 100 KB, and genes with ≥2 enhancers within 100 KB were further calculated.

To evaluate the overall enhancing effects of enhancers with different repetitive types, 7323 enhancers were divided into 11 categories (1 category of no repeats and 10 categories of repetitive sequences). The FPKM values of genes within 100 KB of the 11 categories of enhancers were calculated. The categories with obviously high expression levels were further divided by the base pair number and composition.

### Experimental validation by dual-luciferase reporter assay system

The 7323 identified enhancers were divided into strong (>4-fold enrichment), medium (2 to 4-fold enrichment), and weak (<2-fold enrichment) groups according to their predicted enrichment values [25]. The top 100 enhancers in each group were chosen, of which 15 STARR-seq sites were randomly selected from each group (45 in total). A dual-luciferase reporter assay system was performed on the lower epidermis of tobacco leaves for each enhancer. To fully demonstrate the effects of the enhancers, we modified the original pGreenⅡ0800-*Luc* vector by adding a minimal CaMV 35S promoter before *firefly luciferase*. The modified pGreenⅡ0800-*Luc* was linearized using KpnI, and 45 enhancer sequences were separately ligated into the cutting site before the minimal CaMV 35S promoter. The PCR primers are listed in Table S7.

A CaMV 35S promoter was added before *firefly luciferase* of the original pGreenⅡ0800-*Luc* as a positive control, and it showed strong luciferase fluorescence. Empty modified pGreenⅡ0800-*Luc* was used as a negative control, and it showed almost no luciferase fluorescence. Moreover, the previously reported wheat enhancer *cab-1* was ligated into the modified pGreenⅡ0800-*Luc* as another positive control.After *Agrobacterium* transformation and positive detection, the three control groups were synchronously injected with different parts of the same *N. tabacum L*. leaves, with one STARR-seq enhancer waiting for verification. After 36–48 h of cultivation, D-luciferin was uniformly smeared on the lower epidermis and luciferase fluorescence was imaged using Bio-Rad Image Lab 5.0. The RNA of the tobacco leaves under different treatments was separately extracted using the Plant RNA Collection Kit v1.5 (Catalog No. RN33050, Biofit). RT-qPCR for *luciferase* was performed to quantify the effects of each predicted enhancer. *Renilla luciferase* was used as an internal control.

### Histone modification analysis based on available ChIP-seq data

To reveal the histone modification levels of barley STARR-seq enhancers, ChIP-seq data for barley were downloaded from the NCBI Gene Expression Omnibus (GEO) database (accession number: GSE122539) [86], including data on H3K27me3 and H3K4me3. MACS2 was used to call the peaks of the obtained histone modification data [87], and Bowtie 2 was used to align the sequencing data of both the treatment and control samples to the MorexV2.44_pseudomolecules_assembly genome [88]. deepTools was then utilized to count the number of ChIP-seq reads overlapped in a ±5 KB range from the STARR-seq enhancers, and plot profiles were configured by the “computeMatrix” function in deepTools [89].

## Data availability

The raw STARR-seq data in this study have been deposited in the Genome Sequence Archive [90] in National Genomics Data Center [91], China National Center for Bioinformation / Beijing Institute of Genomics, Chinese Academy of Sciences (GSA: CRA009174) that are publicly accessible at https://ngdc.cncb.ac.cn/gsa.

## CRediT author statement

**Wanlin Zhou**: Writing−Original draft preparation, Validation, Formal analysis **Haoran Shi**: Software, Formal analysis, Visualization **Zhiqiang Wang**: Methodology, Investigation **Yuxin Huang**: Validation, Investigation **Lin Ni**: Validation, Investigation **Xudong Chen**: Resources, Data Curation **Yan Liu**: Writing-Reviewing and Editing **Haojie Li**: Validation, Investigation **Caixia Li**: Writing-Reviewing and Editing **Yaxi Liu**: Conceptualization, Supervision, Funding acquisition

## Acknowledgments

We are grateful for the Key Program of Sichuan Province Natural Science Foundation (2022NSFSC0015) and the Key Research and Development Program of Sichuan Province (2021YFN0034 and 2021YFG0028) in financial help of this work. We would like to thank ***Editage*** (www.editage.cn) for English language proofreading and polishing, and thank peer researchers for providing public RNA-seq data.

## Supplementary material

**Figure S1 Comparison between barley ATAC**-**seq and STARR**-**seq peaks**

**A**. Chromosomal distribution of barley ATAC-seq peaks. Chromosomal length was labeled on the left and the scale bar was on the right. Scale values of the corresponding color represented the number of enhancers per MB (enhancer density).

**B**. Number of overlapped peaks between barley ATAC-seq and STARR-seq data.

**C**. TAC-seq, Assay for Transposase-Accessible Chromatin using sequencing.

**Figure S2 Comparison between rice ATAC**-**seq and STARR**-**seq peaks**

**A**. Chromosomal distribution of rice STARR-seq peaks. **B**. Chromosomal distribution of rice ATAC-seq peaks. Chromosomal length was labeled on the left and the scale bar was on the right. Scale values of the corresponding color represented the number of enhancers per 100 KB (enhancer density)

**Figure S3 Chromosomal distribution of maize ATAC**-**seq peaks**

Chromosomal length was labeled on the left and the scale bar was on the right. Scale values of the corresponding color represented the number of enhancers per MB (enhancer density).

**Table S1 Information of identified STARR-seq enhancers in cultivar Morex** Peaks with pValue < 0.001 and enrichment > 1.3 were considered as enhancers; enrichment represented the strength of predicted enhancers.

**Table S2 Information of enhancers in various repeatitive sequence categories via Repeatmasker**

Score, Smith-Waterman score of the match; query sequence was the input enhancer sequence; C/+, C represented match is with the complement of the consensus sequence in the database; + represented match is with the same sequence. Names and class of the matching repeats were listed in column **G** and **H**.

**Table S3 FPKM data of genes within different range of STARR-seq enhancers Supplementary Table3a**: FPKM data of genes within 10 KB of STARR-seq enhancers;

**supplementary Table3b**: FPKM data of genes within 10−50 KB of STARR-seq enhancers;

**supplementary Table3c**: FPKM data of genes within 50−100 KB of STARR-seq enhancers;

**supplementary Table3d**: FPKM data of genes ≥ 100 KB of STARR-seq enhancers. RNA-seq data of 228 samples were collected from public database. FPKM, Fragments Per Kilobase Million.

**Table S4 FPKM data of genes having 0, 1, and** ≥ **2 enhancers within 100 KB Supplementary Table4a**: FPKM data of genes having no enhancer within 100 KB; **Supplementary Table4b**: FPKM data of genes having 1 enhancer within 100 KB;

**Supplementary Table4c**: FPKM data of genes having = or > 2 enhancers within 100 KB. RNA-seq data of 228 samples were collected from public database.

**Table S5 FPKM data of genes within 100 KB of enhancers in 11 different repeat categories**

**Supplementary Table5a**: FPKM data of barley genes within 100 KB of “DNA/TcMar-Stowaw” enhancers;**Supplementary Table5b:** FPKM data of barley genes within 100 KB of “tRNA” enhancers;**Supplementary Table5c:** FPKM data of barley genes within 100 KB of “DNA/MULE-MuDR” enhancers;**Supplementary Table5d:** FPKM data of barley genes within 100 KB of “satellite” enhancers;

**Supplementary Table5e:** FPKM data of barley genes within 100 KB of “LINE/L1” enhancers;**Supplementary Table5f:** FPKM data of barley genes within 100 KB of “low-complexity” enhancers;**Supplementary Table5g:** FPKM data of barley genes within 100 KB of “DNA CMC-EnSpm” enhancers;**Supplementary Table5h:** FPKM data of barley genes within 100 KB of “simple repeat” enhancers;**Supplementary Table5i:** FPKM data of barley genes within 100 KB of “LTR/Copia” enhancers;**Supplementary Table5j:** FPKM data of barley genes within 100 KB of “LTR/Gypsy” enhancers;**Supplementary Table5k:** FPKM data of barley genes within 100 KB of no repetitive enhancers. RNA-seq data of 228 samples were collected from public database.

**Table S6 FPKM data of genes within 100 KB of enhancers with different repetitive base pair composition Supplementary Table6a:** FPKM data of barley genes within 100 KB of “satellite” enhancers;**Supplementary Table6b:** FPKM data of barley genes within 100 KB of 1-bp repeat enhancers;**Supplementary Table6c:** FPKM data of barley genes within 100 KB of 2-bp repeats enhancers;**Supplementary Table6d:** FPKM data of barley genes within 100 KB of 3-bp repeats enhancers;**Supplementary Table6e:** FPKM data of barley genes within 100 KB of 4-bp repeats enhancers;**Supplementary Table6f:** FPKM data of barley genes within 100 KB of 5-bp repeats enhancers;**Supplementary Table6g:** FPKM data of barley genes within 100 KB of 6-bp repeats enhancers;**Supplementary Table6h:** FPKM data of barley genes within 100 KB of 2-bp repeats enhancers with different base pair composition;**Supplementary Table6i:** FPKM data of barley genes within 100 KB of 6-bp repeats enhancers with different base pair composition. RNA-seq data of 228 samples were collected from public database.

**Table S7 Primers used in enhancers’ vector construction** Se, Strong enhancer; Me, Medium enhancer; We, Weak enhancer.

