## Supplementary figures and images for "Identification of Barley Enhancers across Genome via STARR-seq"

### Figure S1

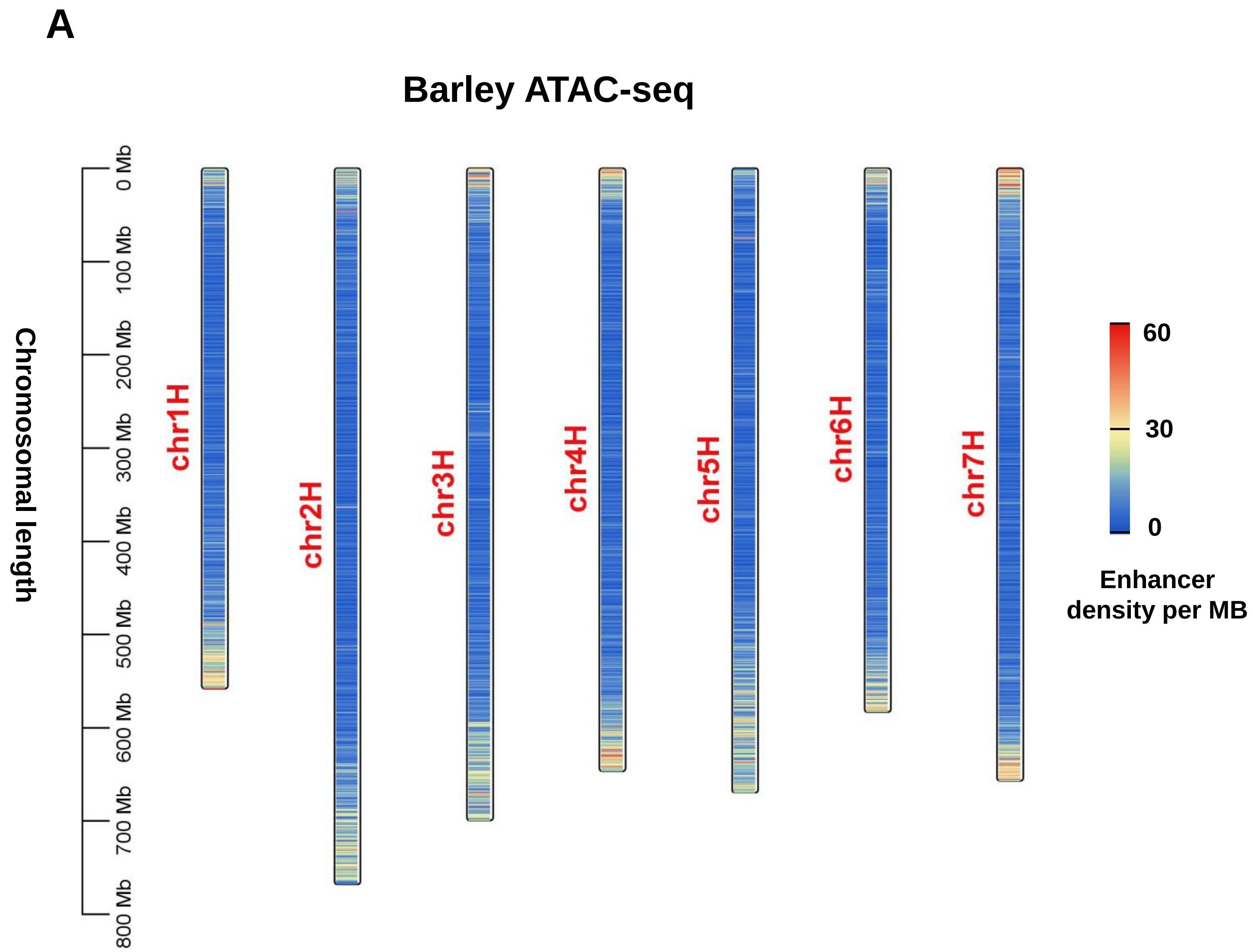

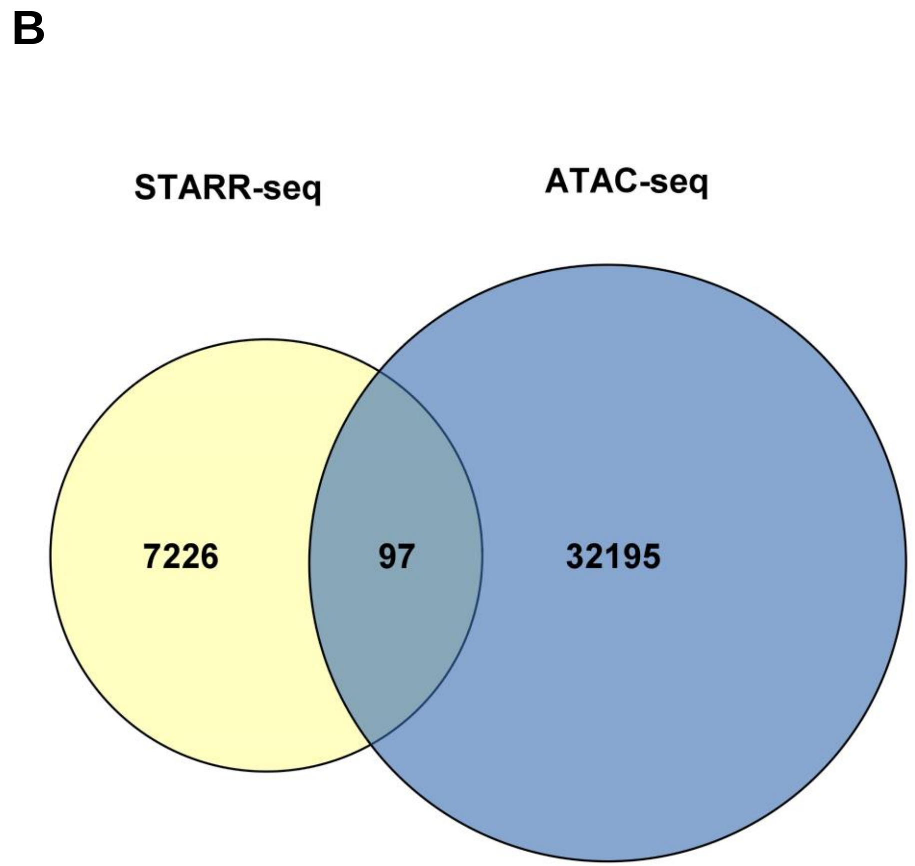

### Figure S2

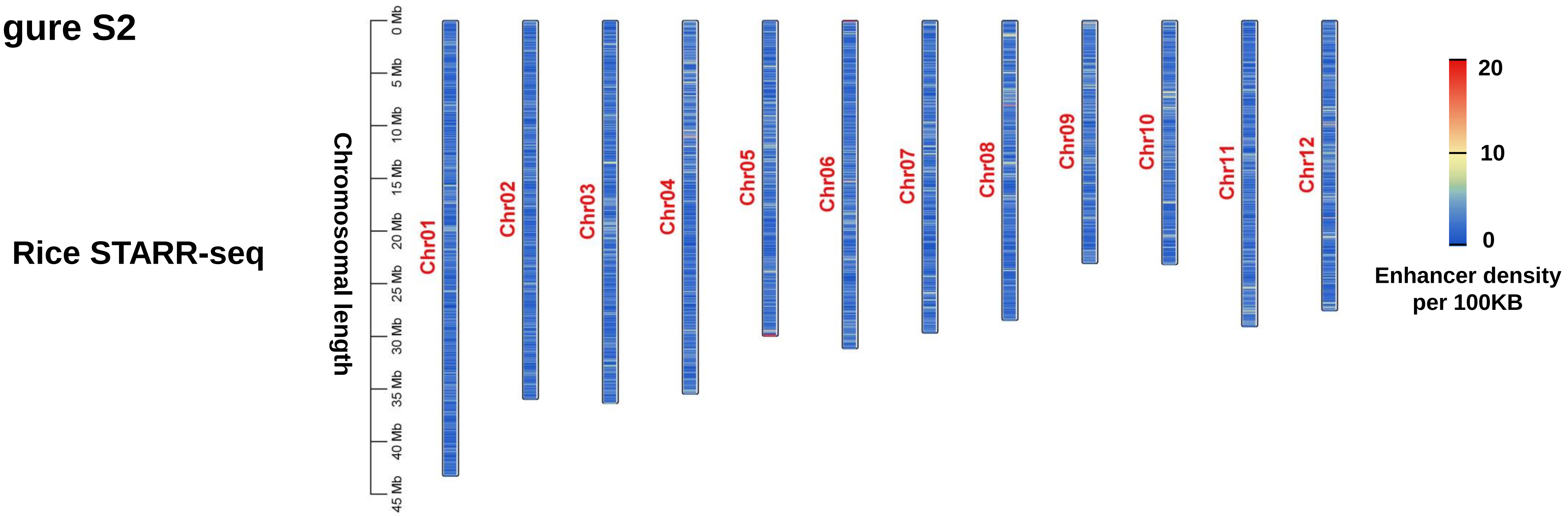

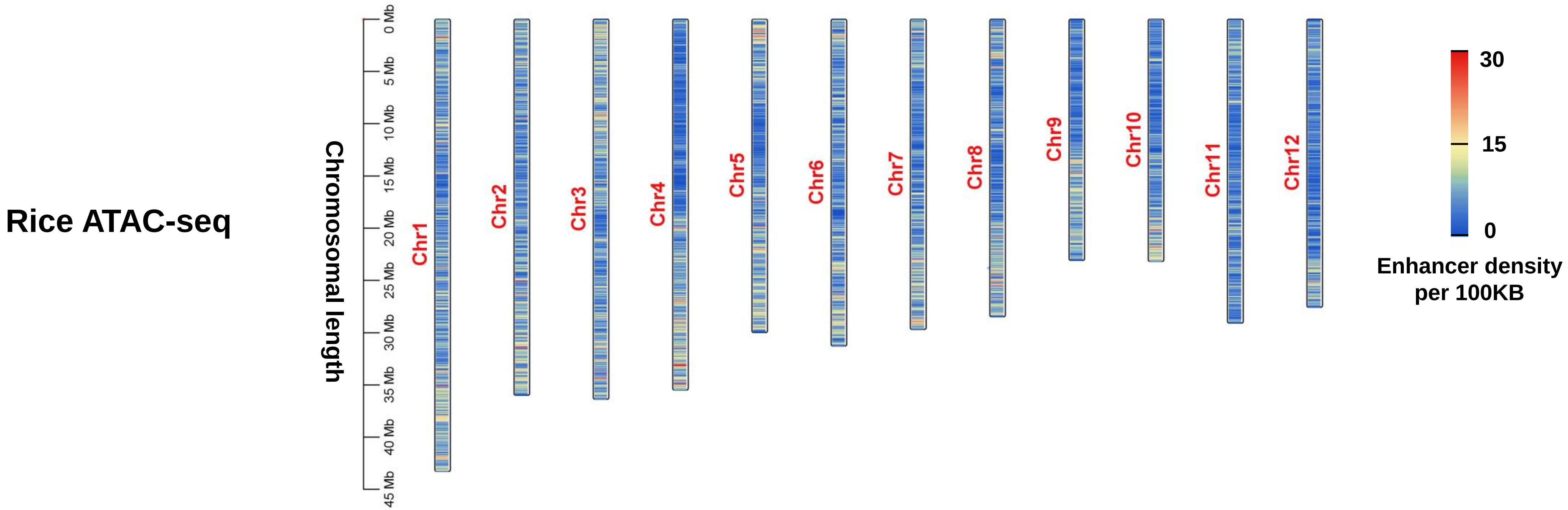

### Figure S3

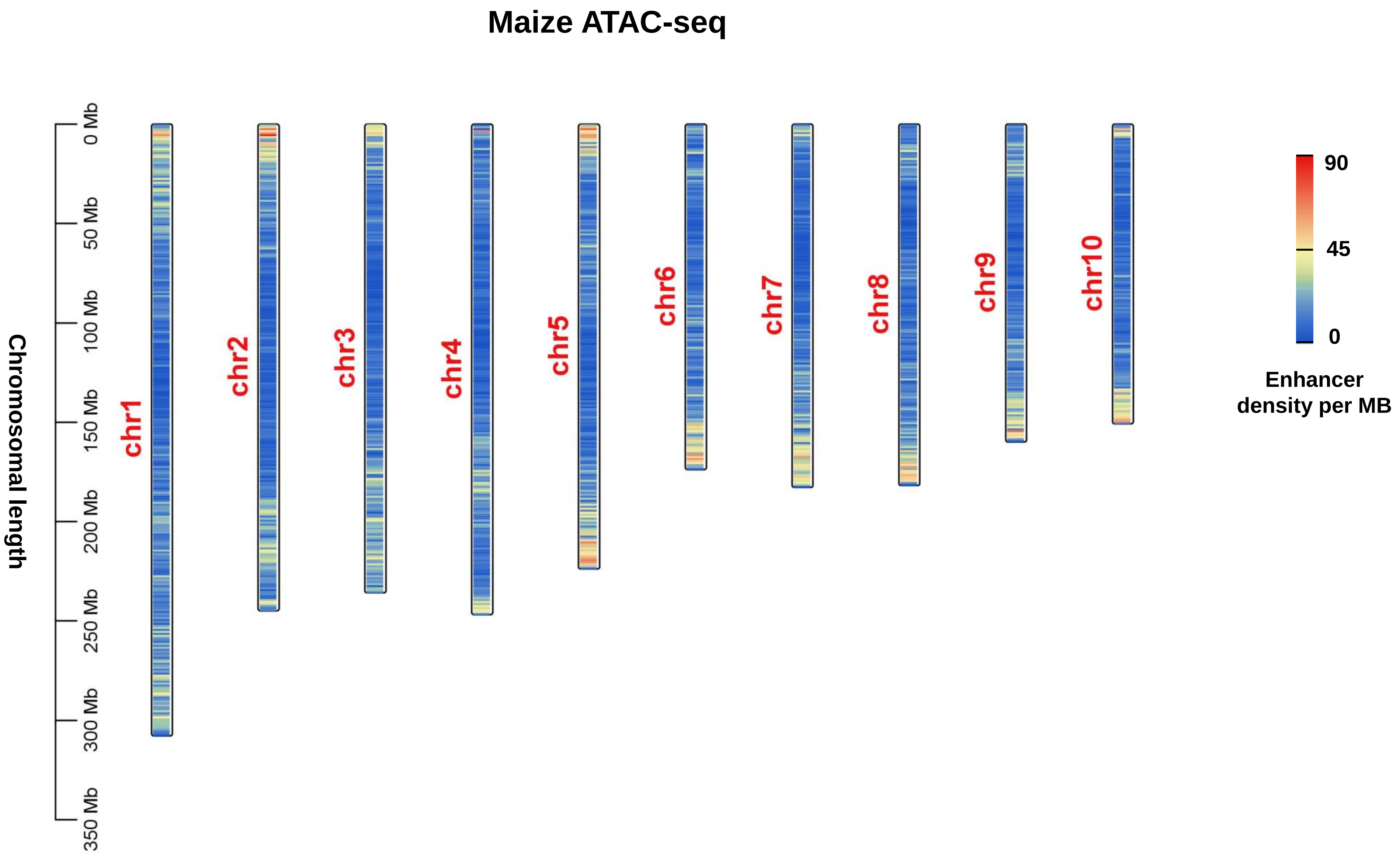
